# Threonine phosphorylation regulates the molecular assembly and signaling of EGFR in cooperation with membrane lipids

**DOI:** 10.1101/2021.05.14.444132

**Authors:** Ryo Maeda, Hiroko Tamagaki-Asahina, Takeshi Sato, Masataka Yanagawa, Yasushi Sako

**Affiliations:** Cellular Informatics Laboratory, RIKEN CPR, Wako, Saitama 351-0198, Japan; Kyoto Pharmaceutical University, 5, Misasagi-cho, Yamashina, Kyoto, 607-8414, Japan; CREST JST, 4-1-8, Honcho, Kawaguchi, 332-0012, Japan

## Abstract

The cytoplasmic domain of the receptor tyrosine kinases (RTKs) plays roles as a phosphorylation enzyme and a protein scaffold but the allocation of these two functions is not fully understood. We here analyzed assembly of the transmembrane (TM)-juxtamembrane (JM) region of EGFR, one of the best studied species of RTKs, by combining single-pair FRET imaging and a nanodisc technique. The JM domain of EGFR contains a threonine residue (Thr654) that is phosphorylated after ligand association. We observed that the TM-JM peptides of EGFR form anionic lipid-induced dimers and cholesterol-induced oligomers. The two forms involve distinct molecular interactions, with a bias towards oligomer formation upon threonine phosphorylation. We further analyzed the functions and oligomerization of whole EGFR molecules, with or without a substitution of Thr654 to alanine, in living cells. The results suggested an autoregulatory mechanism in which Thr654 phosphorylation causes a switch of the major function of EGFR from kinase activation dimers to scaffolding oligomers.

## Introduction

Epidermal growth factor receptor (EGFR) is an RTK responsible for cell proliferation and differentiation (1, 2) and consists of five domains; an extracellular domain that interacts with extracellular ligands, a single-pass transmembrane (TM) helix, a juxtamembrane (JM) domain, a cytoplasmic kinase domain, and a C-terminal tail domain for interaction with various cytoplasmic proteins (3, 4). Ligand association changes the conformation of EGFR in its extracellular domain (5) and induces formation of an asymmetric dimer of the intracellular kinase domains (6). This dimerization subsequently results in the phosphorylation of tyrosine residues on the tail domain and the recruitment of intracellular signal proteins such as GRB2 and PLCγ containing SH2 and/or PTB domains (7). Although the atomic structures of most of the EGFR domains excluding the tail domain have been elucidated individually (5, 6, 8–10), the overall architecture of this protein has not yet been revealed, leaving several unanswered questions about the molecular mechanisms underlying its functions. The correlation between the arrangement of EGFR molecules and their function is therefore still controversial, e.g., it has long been established that the dimerization of EGFR is necessary and sufficient for kinase activation (11), whereas several studies have reported the importance of higher-order oligomerization for EGFR-mediated signal transduction (12–14).

The TM helix and the JM domain (TM-JM) of EGFR play important roles in the conformational coupling of ligand binding to its activation and oligomerization (9, 11, 15). Previous NMR studies and molecular dynamics simulations have suggested that the TM domain forms an α-helix dimer that undergoes a change in the interaction between GXXXG motives (9, 16, 17) following the ligand association with its extracellular domains (17, 18). This information regarding conformational changes in the TM dimer is then transmitted to the JM domain which comprises a JM-A (N-terminal half) region that can form an antiparallel helix dimer, and a JM-B (C-terminal half) region which makes intramolecular contact with the kinase domain (11). Both these JM regions contribute to the stable formation of an asymmetric kinase dimer conformation, which is crucial for kinase activation. The JM-A domain is rich in Lys and Arg residues, several of which are thought to interact with anionic lipid molecules of the plasma membrane and promote antiparallel dimer formation (19–21). In addition to the phospholipid species, cholesterol is a major component of the plasma membrane, mainly distributed as lipid rafts and caveolae, and has been implicated in the regulation of membrane fluidity and receptor function. Previous studies have shown that EGFR molecules are clustered in lipid rafts (14, 22), suggesting an interaction with cholesterols. Of note in this regard, it has been reported that the depletion of cholesterols induces various effects on EGFR signaling, also this remains controversial (23–25). Another important factor in the regulation of EGFR through the TM-JM is the phosphorylation of Thr654 at the JM-A domain. Although Thr phosphorylation is known to be involved in EGFR deactivation, the precise mechanism of this is still elusive (26).

In our present study, by combining single-pair FRET measurements and nanodisc technology, we studied how the functions of anionic lipids, cholesterols, and EGFR Thr654 phosphorylation (pT654) are orchestrated to achieve the regulation of dimerization and/or oligomerization of EGFR. We previously reported that anionic lipids cause the dimerization of JM domains, and that pT654 together with acidic lipids induces the dissociation of the EGFR dimer (20). In this current study, we report that both the TM and JM protomers of EGFR are positioned closer to each other in the presence of cholesterols than in the EGFR dimers promoted by anionic lipids. Furthermore, we found that TM-JM peptides were oligomerized in cholesterol containing membranes, which was promoted by pT654. Finally, in living cells expressing whole EGFR molecules, we observed differential functional roles and oligomerization states that are dependent on the pT654 levels. Our recent study also shows that EGF-induced oligomerization of EGFR depends on the cholesterol density in the plasma membrane (27). These results suggest that the membrane cholesterol and pT654 cooperatively shift the EGFR function from the kinase dimer to the scaffold oligomer.

## Results

### Incorporation of TM-JM peptides into nanodiscs

Synthesized peptides of the TM-JM region of EGFR were prepared and labeled with a fluorophore Cy3 or Cy5 at the N-terminus (TM terminal region) or C-terminus (JM terminal region), respectively (Fig. 1a). The peptides were reconstituted into nanodisc structures with membrane scaffold proteins (MSPs) and lipid molecules (Fig. 1b, c). Mixtures of POPC (PC), POPS (PS), and cholesterol were used for reconstruction (Fig. 1d). The nanodiscs containing cholesterol showed two peaks following size exclusion gel chromatography, one of which had a smaller disc size relative to that without cholesterol (Fig. 1e). To avoid the effects of disc size, we collected and used nanodiscs involved in the first peak fraction which had a similar size without cholesterol. This fraction was composed of the same ratio of lipids as used in the preparation (Fig. 1f). Fractions of PS (inner leaflet) and cholesterol (inner and outer leaflets) mimicked those in the plasma membrane. Synthesized TM-JM peptides with pT654 were also reconstituted into nanodiscs. The nanodisc construction was examined under a transmission electron microscope (Fig. 1g). In total, 16 types of nanodiscs were applied to subsequent single-molecule measurements.

**Figure 1.**
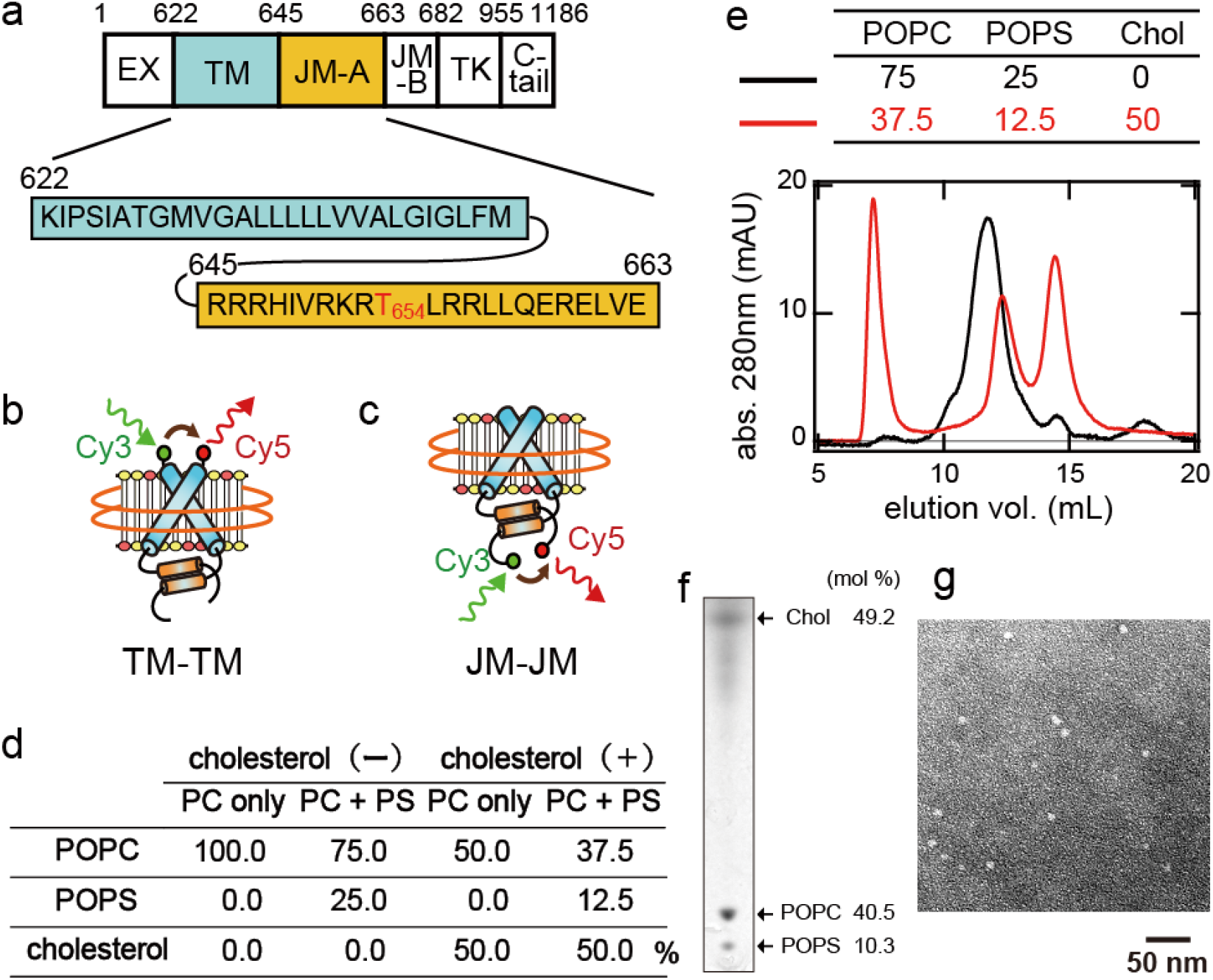
Construction of nanodiscs containing fluorescent TM-JM peptides of EGFR. (**a**) Amino acid sequence of the EGFR TM and JM-A domains. EX, extracellular domain; TK, tyrosine kinase domain. (**b, c**) Schematic images of nanodiscs containing dimeric TM-JM peptides fluorescently labeled at the N- (b) and C-terminus (c), respectively. (**d**) Lipid compositions used in the preparation of each nanodisc sample. The fractional ratios of PS and cholesterol mimic those in the mammalian plasma membranes. (**e**) Size-exclusion chromatography used for the purification of nanodiscs containing cholesterol (red) or not (black). The charge ratios of PC/PS/cholesterol are described in the upper table. The fraction having a peak absorbance of around 12 elution volumes (mL) was collected and used for the subsequent experiments. (**f**) Thin-layer chromatography of the nanodisc fraction containing cholesterol (fraction 12 in (**e)**). (**g**) A negative stain electron micrograph of fraction 12 in **(e)**. Size distribution of the nanodiscs calculated from the images was fitted with a Gaussian function, with a mean diameter of 11 ± 2 nm.

### TM-TM interaction in the EGFR dimer

Nanodiscs containing Cy3 and Cy5-labeled peptides were immobilized onto glass surfaces and illuminated with a 532-nm laser for Cy3 excitation. A portion of the fluorescent spots contained Cy5 fluorescence derived from the occurrence of FRET (Fig. 2a, b). Based on the fluorescence intensity, we selected nanodiscs containing one Cy3- and one Cy5-labeled peptide and calculated the FRET efficiency, *E*_FRET_ (Fig. 2c).

**Figure 2.**
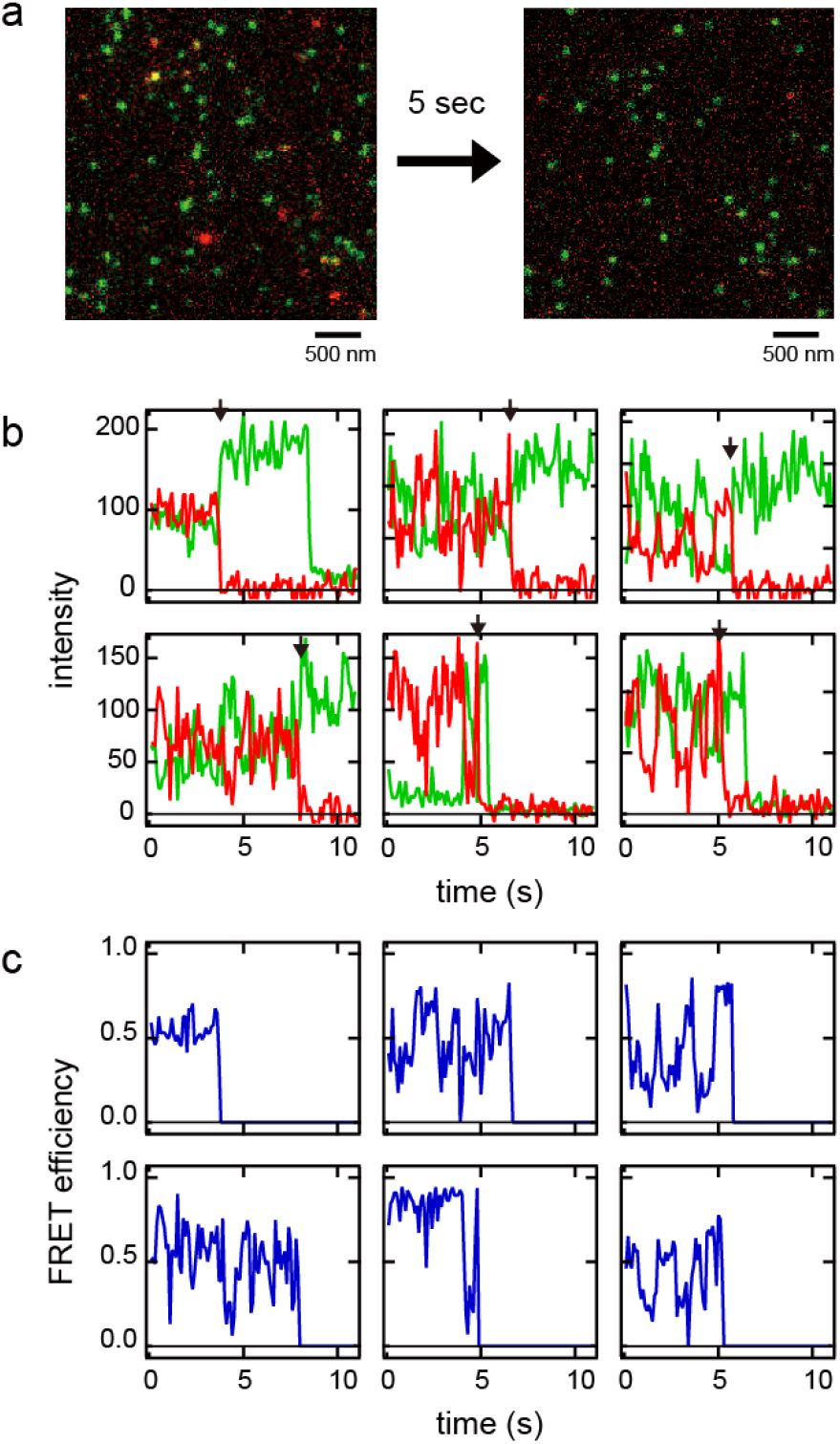
Single-pair FRET measurement of the EGFR TM-JM dimers in nanodiscs. (**a**) Fluorescence micrograph of nanodiscs illuminated with a green laser. Cy3 (green) and Cy5 (red) emissions were superimposed. The Cy5 emission was caused by FRET from Cy3. (**b**) Representative fluorescence trajectories of Cy3 (green) and Cy5 (red). Black arrows indicate photobleaching points of Cy5. (**c**) FRET efficiency trajectories of the fluorescence trajectories in **(b)**. The FRET efficiency, *E*_FRET_, was calculated as described in the Materials and Methods section. Typical fluorescence and FRET trajectories between peptides labeled at the C-terminus are shown. Transitions to low FRET efficiency states suggested that dissociation of the JM dimer occurred occasionally. The Förster radius *R*_0_ between Cy3 and Cy5 is 5.6 nm.

We first examined the interactions between the N-terminal regions of the TM domains (Fig. 3). When Thr654 was not phosphorylated and the membrane contained only PC as lipid species, *E_FRET_* distributed with a peak at a relatively high (> 0.90; Table I) value (Fig. 3a), indicating close proximity of the two TM domains. There may be additional stable structures between the TM domains, as suggested by the small peaks and shoulders in the *E_FRET_* distribution. The addition of anionic lipid PS caused few effects, i.e., the TM dimers were maintained as the major structure (Fig. 3b). Peptides with pT654 slightly decreased the major peak positions of the *E_FRET_* distributions in the PC or PC/PS membranes (Fig. 3e, f). The smooth distribution of the pT654 peptides suggested that pThr654 had homogenized possible substructures of the TM dimers of non-phosphorylated peptides. PS had little effect on the TM-TM interactions regardless of the Thr654 phosphorylation. The presence of cholesterol in the membrane concentrated the distributions to a high *E_FRET_* (> 0.95) region (Fig. 3c, g, h), indicating that the N-terminal regions of TM-TM dimers were positioned in extremely close proximity. Although the peak position was similar in PC and PC/cholesterol membranes (Fig 3a, c; Table I), the distribution in PC/cholesterol membrane was more accumulated at the peak position. It also should be noted that the accumulation at a high *E_FRET_* region was a remarkable observation for pT654 peptides. Thus, pT654 and the presence of membrane cholesterol decreased the distance between the two N-termini of the TM domains in cooperation. PS competed with cholesterol when Thr654 was non-phosphorylated (Fig. 3d).

**Figure 3.**
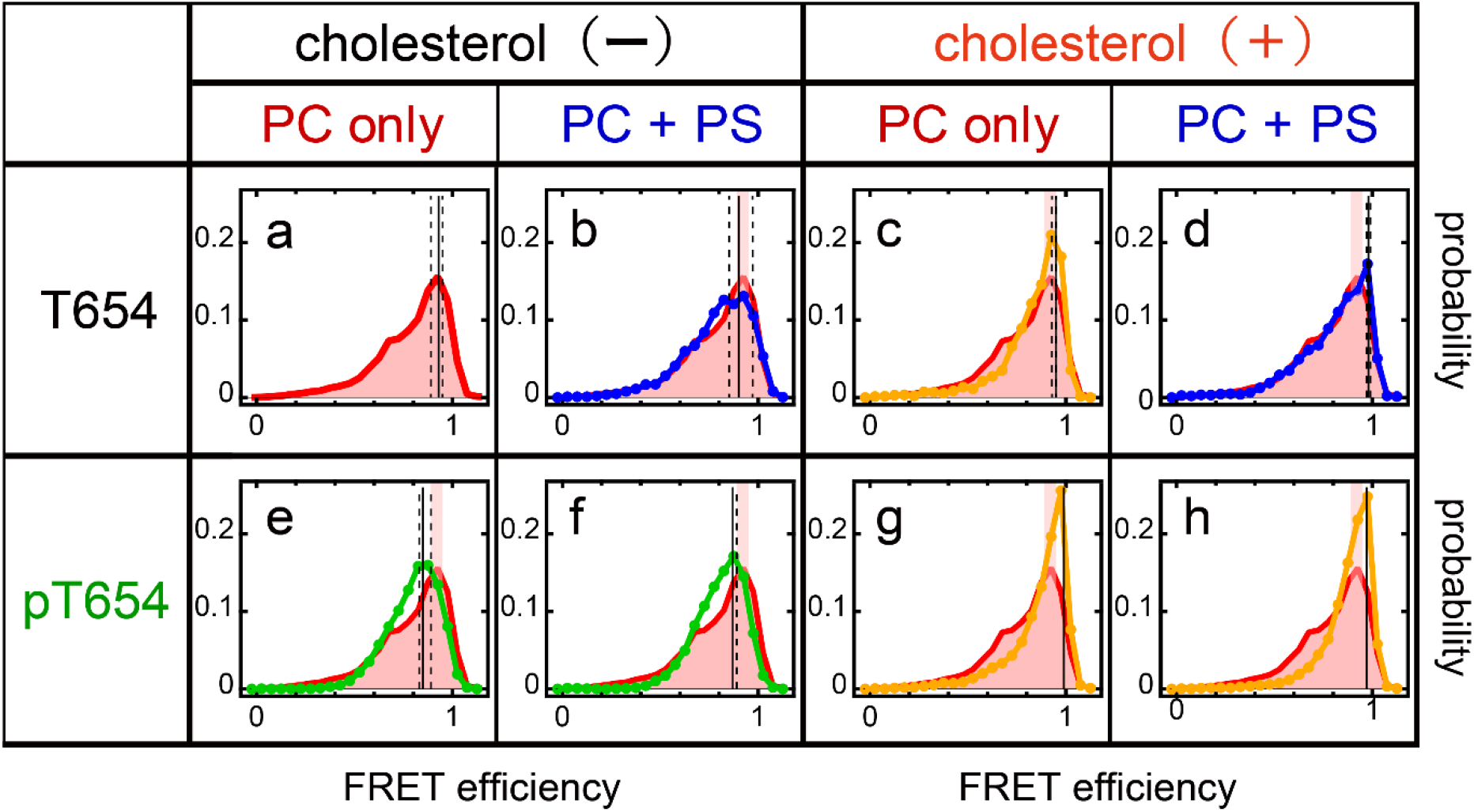
FRET efficiency(*E*_FRET_) distributions in nanodiscs containing a single Cy3/Cy5-pair of N-terminal labeled peptides. Nanodiscs contained non-phosphorylated (**a–d**) and Thr654 phosphorylated (**e–h**) peptides at the indicated lipid conditions. Positions of the mode and its 5-95% percentile section are indicated by solid and dashed lines, respectively. See Table I for these values. In **(b–h)**, the distribution and the 95% percentile section shown in **(a)** are superimposed (red) for comparison.

**Table I.** Peak values in the *E_FRET_* distributions between the TM-JM peptides.

| lipid composition | pT654 | labelling |  |  |  |
| --- | --- | --- | --- | --- | --- |
|  |  | N-term |  | C-term |  |
|  |  | mean | 95% | mean | 95% |
| PC | - | 0.93 | (0.89-0.95) | 0.81 | (0.73-0.87) |
| PC/PS | - | 0.90 | (0.85-0.97) | 0.83 | (0.79-0.89) |
| PC/chol | - | 0.95 | (0.93-0.95) | 0.93 | (0.85-0.97) |
| PC/PS/chol | - | 0.98 | (0.97-0.99) | 0.94 | (0.91-0.97) |
| PC | + | 0.85 | (0.83-0.89) | 0.68 | (0.63-0.89) |
| PC/PS | + | 0.87 | (0.87-0.89) | 0.36 | (0.33-0.43) |
| PC/chol | + | 0.99 | (0.99) | 0.90 | (0.89-0.95) |
| PC/PS/chol | + | 0.97 | (0.97) | 0.82 | (0.81-0.89) |

### JM-JM interaction in the EGFR dimer

To examine the effects of lipid species and pT654 on JM-JM interaction, *E_FRET_* distributions were determined under the C-terminus labeling (Fig. 4). In PC membrane, *E_FRET_* was broadly distributed with a peak around 0.81 (Fig. 4a; Table I). It is plausible that the JM-A dimers are fluctuating between minor dissociation and major association states. In the PC/PS membrane, the peak fraction was increased, indicating that PS stabilized the JM-A dimer conformation (Fig. 4b). pT654 increased the low FRET fraction in the PC/PS membrane (Fig. 4f) but showed little effect in the PC membrane (Fig. 4e). These results confirmed the results of our previous study (20). Cholesterol moved the *E_FRET_* peak between non-phosphorylated peptides to higher values (0.93–0.94) regardless of whether it was a PC or PC/PS membrane (Fig. 4c, d), i.e., cholesterol forced the C-termini of the JM-A domains to position closer.

**Figure 4.**
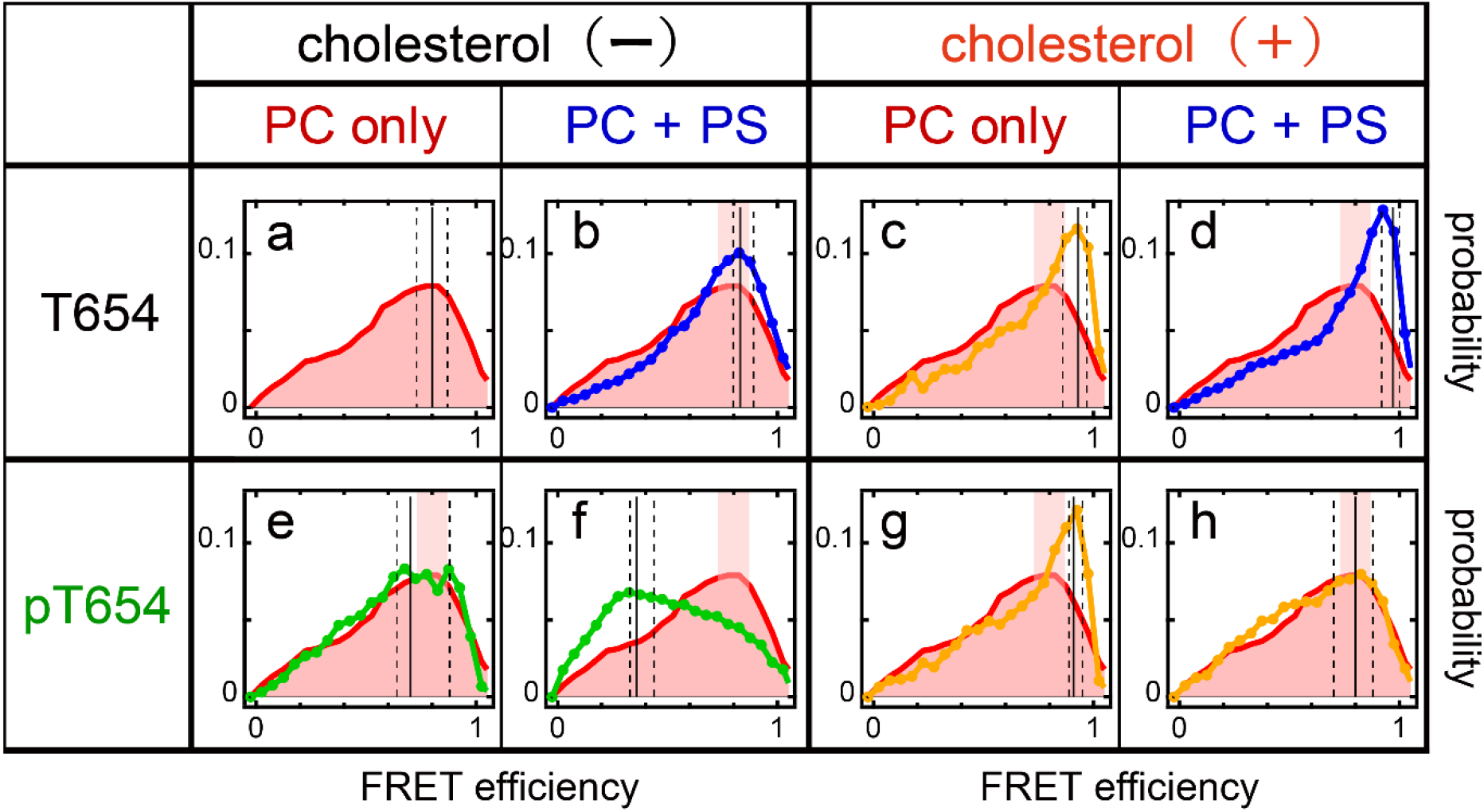
*E*_FRET_ distributions in nanodiscs containing a single Cy3/Cy5-pair of C-terminal labeled peptides. Nanodiscs contained non-phosphorylated (**a–d**) and Thr654 phosphorylated (**e–h**) peptides in the indicated lipid conditions. Positions of the mode and its 5-95% percentile section are indicated by solid and dashed lines, respectively. See Table I for these values. In **(b–h)**, the distribution and the 95% percentile section shown in **(a)** are superimposed (red) for comparison.

The mixed effects of cholesterol and pT654 on the JM-JM interaction were further examined. In the PC membrane (Fig. 4g), cholesterol increased the high FRET population to a comparable level to those shown for non-phosphorylated peptides. Cholesterol in the PC/PS membrane (Fig. 4h) reversed the *E_FRET_* distribution seen in the PC/PS membrane without cholesterol (Fig. 4f) to that observed in the PC membrane (Fig. 4e) i.e. minor low FRET and major high FRET states in the PC/PS/cholesterol membrane. However, it should be noted that the *E_FRET_* values were not as large as those found in other conditions with cholesterol (Fig. 4c, d, g), i.e., the cholesterol effect on JM-JM interaction was partially diminished by the coexistence of PS and pThr654. Overall, our data showed that cholesterol increased the proximity between the C-terminus of JM domains in PC and PC/PS membranes, and that this effect overrode that of pThr654 in the PC/PS membrane.

### Higher-order oligomerization of TM-JM peptides

We speculated that the accumulation of EGFR in lipid rafts, which has been reported in previous studies, could be an effect of cholesterol in the raft membrane and examined the assembly of TM-JM peptides in the nanodiscs, collecting images of fluorescent spots containing only Cy3-labeled peptides to avoid interference from the effects of FRET occurring between Cy3 and Cy5. Figure 5 displays the fluorescence intensity histograms of C-terminus-labeled TM-JM peptides in nanodiscs containing or not-containing cholesterol. Cholesterol shifted the histograms toward higher intensities for the pT654 peptides (Fig. 5b, d), suggesting a cooperative effect of cholesterol and Thr phosphorylation to induce higher-order assembly of the TM-JM peptides.

**Figure 5.**
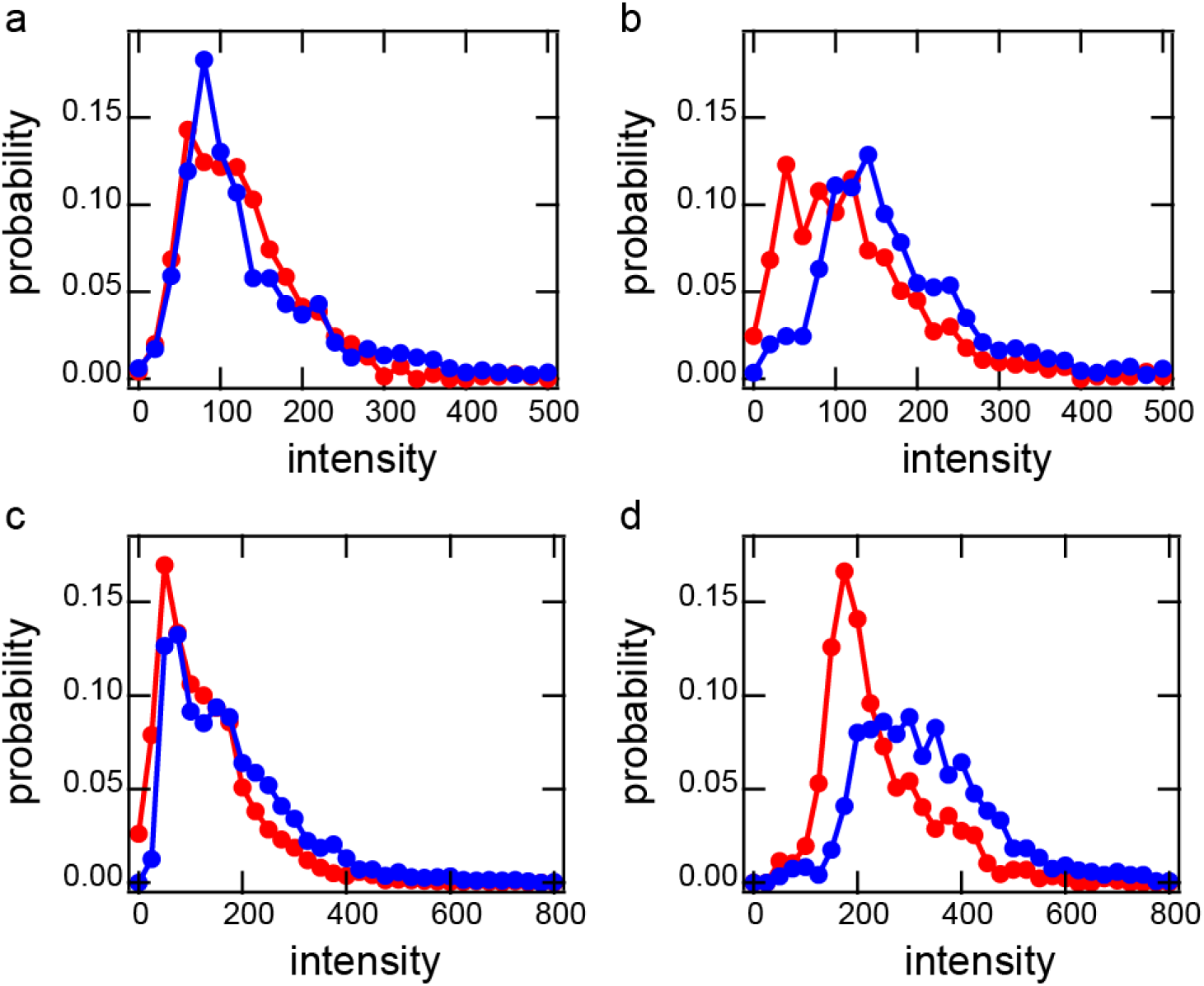
Higher-order oligomerization of EGFR TM-JM peptides in the nanodiscs. Histograms are shown of the total fluorescence intensity of the peptides with Cy3-labeling at the C-terminus in single nanodiscs containing cholesterol (blue) or not (red). Discs containing no Cy5 peptide were chosen for measurement to avoid the possible effects of FRET. Peptides with a non-phosphorylated **(a, c)** or phosphorylated **(b, d)** Thr654 were reconstituted into nanodiscs in PC **(a, b)** or PC/PS **(c, d)** membranes.

### Assembly of TM regions

For analysis of the interactions between more than two TM or JM domains in the assembled structures at the N-terminus, images of fluorescent spots containing two Cy3-labeld peptides and one Cy5-labeled peptide were collected based on their 2-color fluorescence trajectories (Fig. 6a). These nanodiscs showed a variety of Cy3 donor fluorescence intensities before Cy5 photobleaching indicating that the three peptides interacted diversely. We constructed maximum fluorescence intensity histograms in the Cy3 donor channel before and after Cy5 acceptor photobleaching for inference of the interactions between three TM domains (Fig. 6b–i). In all conditions other than non-phosphorylated peptides in the PC/PS membrane, Cy3 distributions after Cy5 photobleaching (red) peaked at the fluorescence intensity of ~100 (in arbitrary units), which was smaller than that observed for the C-terminal-labeled peptides (~150; Fig. 7b– i). This result must have been caused by homo-FRET (self-quenching) between two N-terminal labeled Cy3 peptides. Together with the very small intensity peaks prior to Cy5 photobleaching (blue), these distributions suggested that TM regions of the three peptides (two of them were randomly labeled with Cy3) were oligomerized in very close proximity to each other in the major configuration (trimer; Fig. 6j).

**Figure 6.**
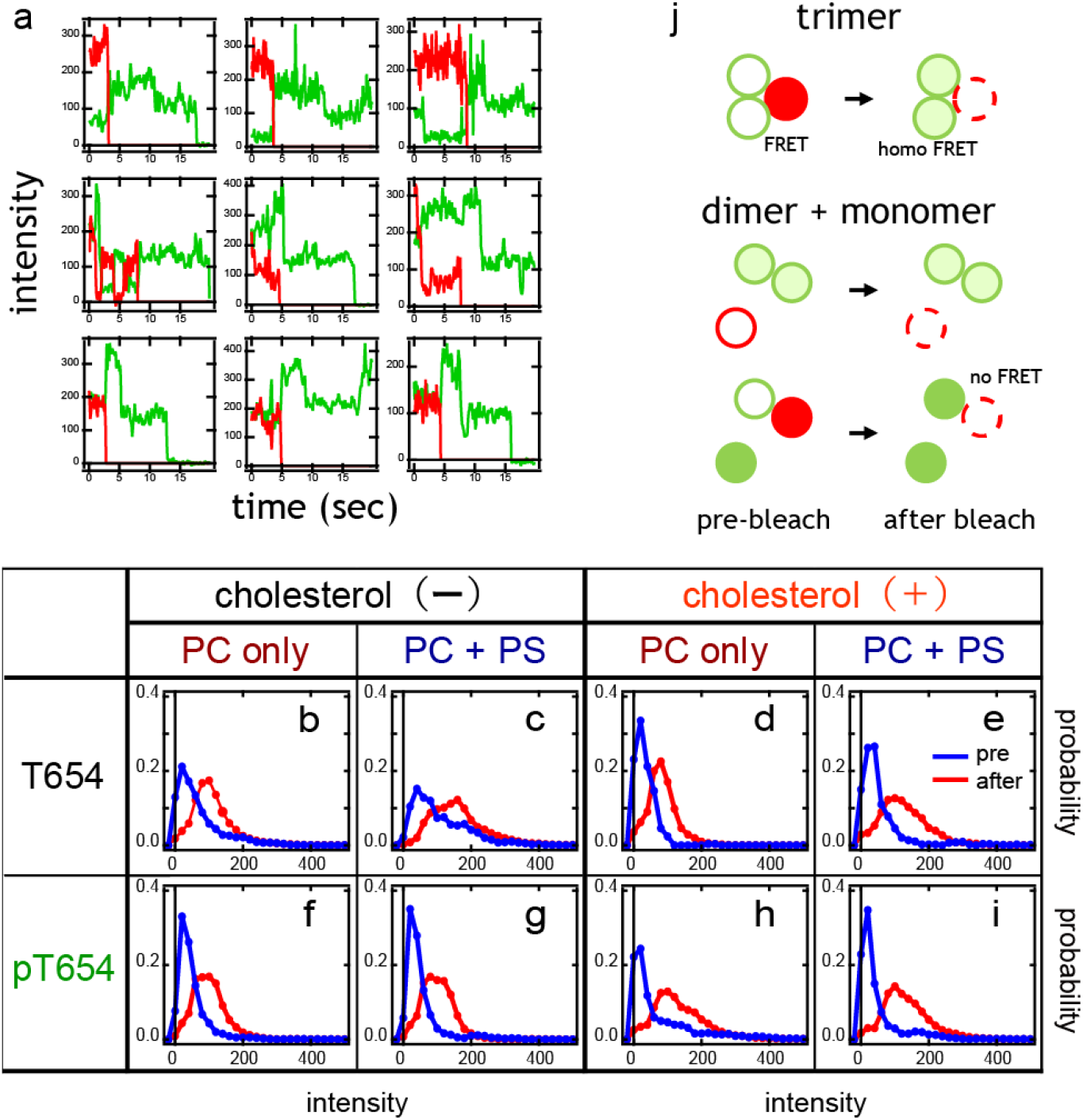
Assembly of EGFR TM regions. (**a**) Representative fluorescence trajectories of Cy3 (green) and Cy5 (red) in nanodiscs containing two Cy3-labeled and one Cy5-labeled peptide. Fluorescence intensities and/or two-step photobleaching dynamics after Cy5 photobleaching indicated that these nanodiscs contained two Cy3 peptides. (**b–i**) Fluorescence intensity histograms of N-terminal-labeled Cy3 peptides before (blue) and after (red) Cy5 photobleaching. Nanodiscs contained non-phosphorylated **(b–e)** and Thr654 phosphorylated **(f–i)** peptides at the indicated lipid conditions. (**j)** Schematic structures indicating proximity between three TM domains before (left) and after (right) Cy5 photobleaching. Note that acceptance of the excitation energy from Cy3 was not saturated for Cy5 under our experimental conditions, even in the presence of two Cy3 molecules.

**Figure 7.**
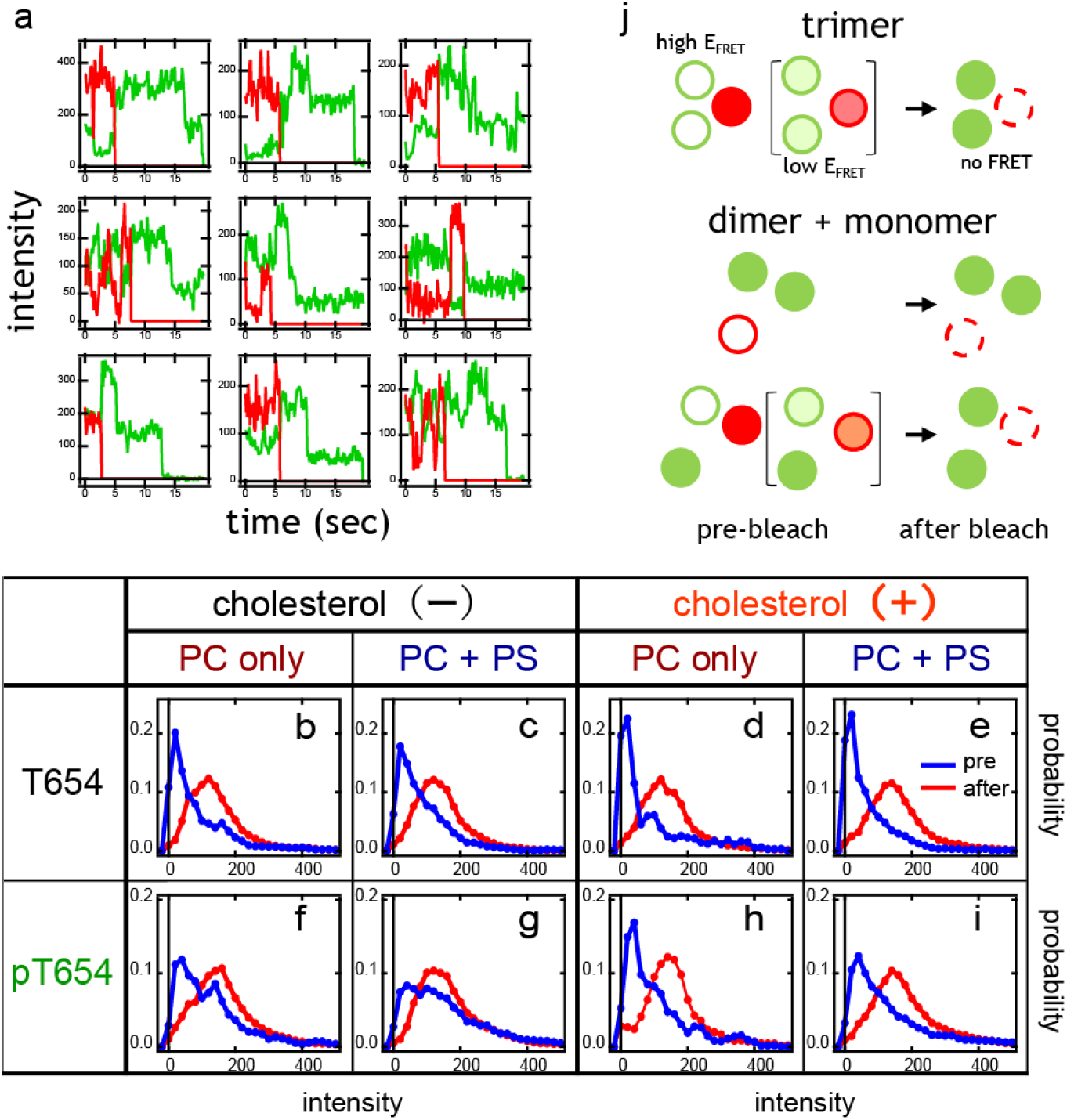
Assembly of EGFR JM regions. (**a**) Representative fluorescence trajectories of Cy3 (green) and Cy5 (red) in nanodiscs containing two Cy3-labeled and one Cy5-labeled peptide at the C-terminus. (**b–i**) Fluorescence intensity histograms of C-terminal-labeled Cy3 peptides from nanodiscs containing two Cy3 and one Cy5 peptide before (blue) and after (red) Cy5 photobleaching. Nanodiscs contained non-phosphorylated **(b–e)** and Thr654 phosphorylated **(f–i)** peptides at the indicated lipid conditions. (**j)** Schematic structures indicating proximity between three JM domains before (left and middle) and after (right) Cy5 photobleaching.

For non-phosphorylated peptides in the PC/PS membrane however (Fig. 6c), the Cy3 intensity histogram after Cy5 photobleaching (red) had a peak intensity at ~150, indicating that two Cy3-labeled peptides in the major population were positioned separately. In addition, the low intensity shoulder in this distribution indicated the presence of proximate dimers (and trimers). Taken together, these distributions suggested that N-terminal regions of three non-phosphorylated peptides have a stronger tendency to arrange as one dimer and one monomer in the PC/PS membrane than any other condition. A similar dimer + monomer arrangement might be contained in the distributions under other conditions as a minor fraction. Consistent with this suggestion, for the non-phosphorylated peptides in PC/PS membrane before Cy5 photobleaching (Fig. 6c, blue), a homo-FRET fraction (~100; with a low *E_FRET_* to Cy5) was evident compared to other conditions. It should be noted that the ability of PS to promote the dimer + monomer arrangement was diminished for pT654 peptides (Fig. 6g). Cholesterol also reduced this effect of PS even for non-phosphorylated peptides (Fig. 6e). As observed in the earlier analysis of the TM-JM peptide dimer (Fig. 3), the effects of PS and cholesterol were competitive.

### Assembly of JM regions

We constructed Cy3 fluorescence intensity histograms of two Cy3 and one Cy5 peptide with C-terminal-labeling in single nanodiscs in order to analyze the interactions between three JM domains (Fig. 7). The distributions of the Cy3 florescence after Cy5 photobleaching (red) were similar under all conditions, exhibiting a single peak at ~150, which was the fluorescence intensity of the two Cy3 molecules without strong interactions to induce homo-FRET. Both the trimer and dimer + monomer arrangements are possible if we assume that the three molecules in the trimer and two molecules in the dimer are not so close that they will induce homo FRET (Fig. 7j).

Prior to Cy5 photobleaching (blue), the distribution peaks were observed in the region of small Cy3 intensities indicating the proximity of both Cy3 molecules with Cy5 to induce high *E_FRET_*, as observed between two molecules in a nanodisc (Fig. 4), i.e., the formation of a JM trimer. An accumulation in the low intensity peak fraction was very evident for non-phosphorylated peptides in the membranes containing cholesterol (Fig. 7d, e). On the other hand, fractions at the intensities similar to those observed after Cy5 photobleaching were significant for pT654 peptides in the membrane without cholesterol (Fig. 7f, g). In general, pT654 peptides exhibited higher fluorescence intensity compared to non-phosphorylated peptides in the corresponding membrane lipid compositions. One possible explanation is that the fraction of high Cy3 intensity before Cy5 photobleaching represents a Cy3 dimer in the dimer + monomer arrangement of three peptides (Fig. 7j). Another possibility is that it was caused by an increased distance between three JM domains in trimers, resulting from Thr phosphorylation to reduce *E_FRET_* (Fig. 7j top middle). These two arrangements could potentially coexist.

Considering the possible arrangement for the TM and JM regions of three TM-JM peptides together (Figs. 6 and 7), we conclude that cholesterol induces the closely proximate oligomerization of three JM domains of non-phosphorylated peptides whereas PS preferentially causes a dimer + monomer arrangement, and the Thr phosphorylation disrupts the JM dimer and facilitates oligomerization of peptides with separated JM domains in the presence of cholesterols (Fig. 8). Because of the limitation of FRET measurement, we could not distinguish whether the peptides in the oligomers directly interacted or not.

**Figure 8.**
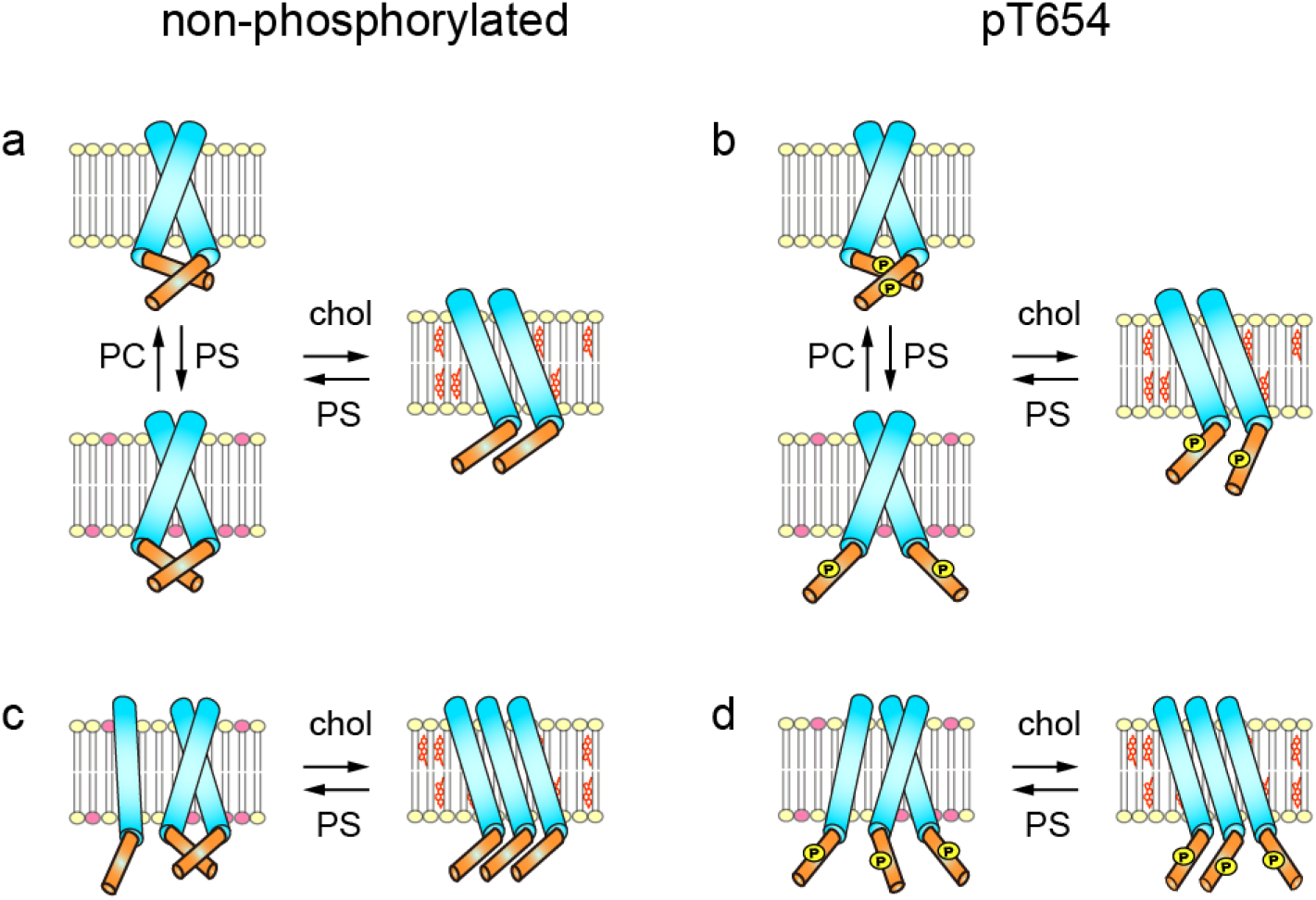
Possible configurations of EGFR TM-JM dimers and trimers regulated by membrane lipids and Thr654 phosphorylation.

### Effect of Thr phosphorylation on the Tyr phosphorylation of EGFR

Our single-molecule structural analysis suggested that pT654 is a key regulator of the molecular assembly of EGFR, which may affect its functions. We examined this possibility in living cells. It is known that PKC activation under EGF signaling induces pT654 in EGFR. This process has been thought to be a negative feedback pathway in the EGFR system. We expressed a wild type (wt) or T654A mutant EGFR in CHO-K1 cells, which have no intrinsic expression of EGFR. An increase in pT654 was observed for wt EGFR after EGF stimulation, and treatment of these cells with phorbol-12-myristate 13-acetate (PMA), a PKC activator, caused stronger phosphorylation of Thr654 regardless of EGF stimulation (Fig. 9a). Application of a saturation amount (100 ng/ml) of EGF to the culture medium induced phosphorylation of Tyr1068 (pY1068) of both the wt and T654A mutant EGFR proteins (Fig. 9b). pY1068 is a major association site on EGFR for the adaptor protein GRB2 and its levels after EGF stimulation were significantly increased by the T654A mutation compared to wt, as expected from the negative effect of pT654. Whereas, pretreatment with PMA decreased the pY1068 level in both the wt and T654A mutant EGFR (Fig. 9c). The PMA-induced decrease in pY1068 for the wt protein could be a negative effect of increased pT654, but the similar result with the T654A mutant suggests that PMA has indirect effects independent of pT654.

**Figure 9.**
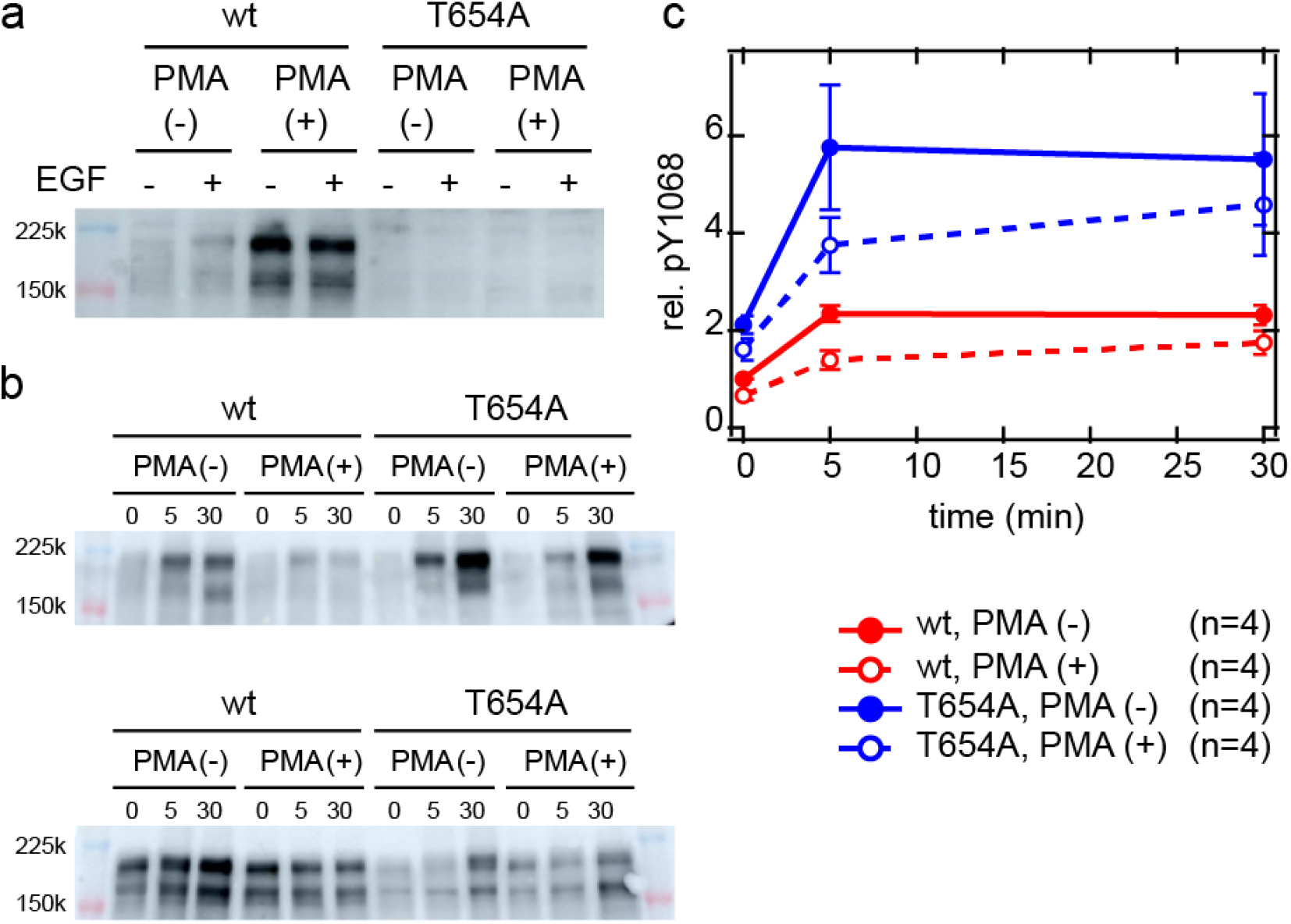
Thr and Tyr phosphorylation of EGFR. (**a**) Thr654 phosphorylation after EGF stimulation and PMA pretreatment. (**b, c**) Timecourses of Y1068 phosphorylation for the wt and T654A mutant of EGFR during EGF stimulation. Typical western blotting results are indicated **(b, top)** and the average of four independent experiments are shown with SE **(c)**. Phosphorylation levels were normalized to the expression levels of the whole EGFR molecule **(b, bottom)**.

### Single-molecule imaging of the clustering and movement of EGFR

We expressed EGFR (wt and T654A) fused with GFP in CHO-K1 cells and, by using single-molecule imaging, detected cluster size distributions and lateral diffusion movements of EGFR molecules in the plasma membrane (Fig. 10a). Clustering of EGFR was measured as the fluorescence intensity distribution of EGFR spots, and the lateral diffusion movements were measured as the increase in the mean square displacement (MSD) of the spots with time. Both measurements were performed before and after 10 min of EGF application to the medium. Application of EGF to the medium induced clustering and immobilization of wt EGFR as we have reported previously(14, 28). The distributions of EGFR cluster size suggest formation of oligomers containing up to more than 10 molecules (Fig. 10b). The convex shapes of MSD curve with time indicate subdiffusion of EGFR molecules (Fig. 10c).

**Figure 10.**
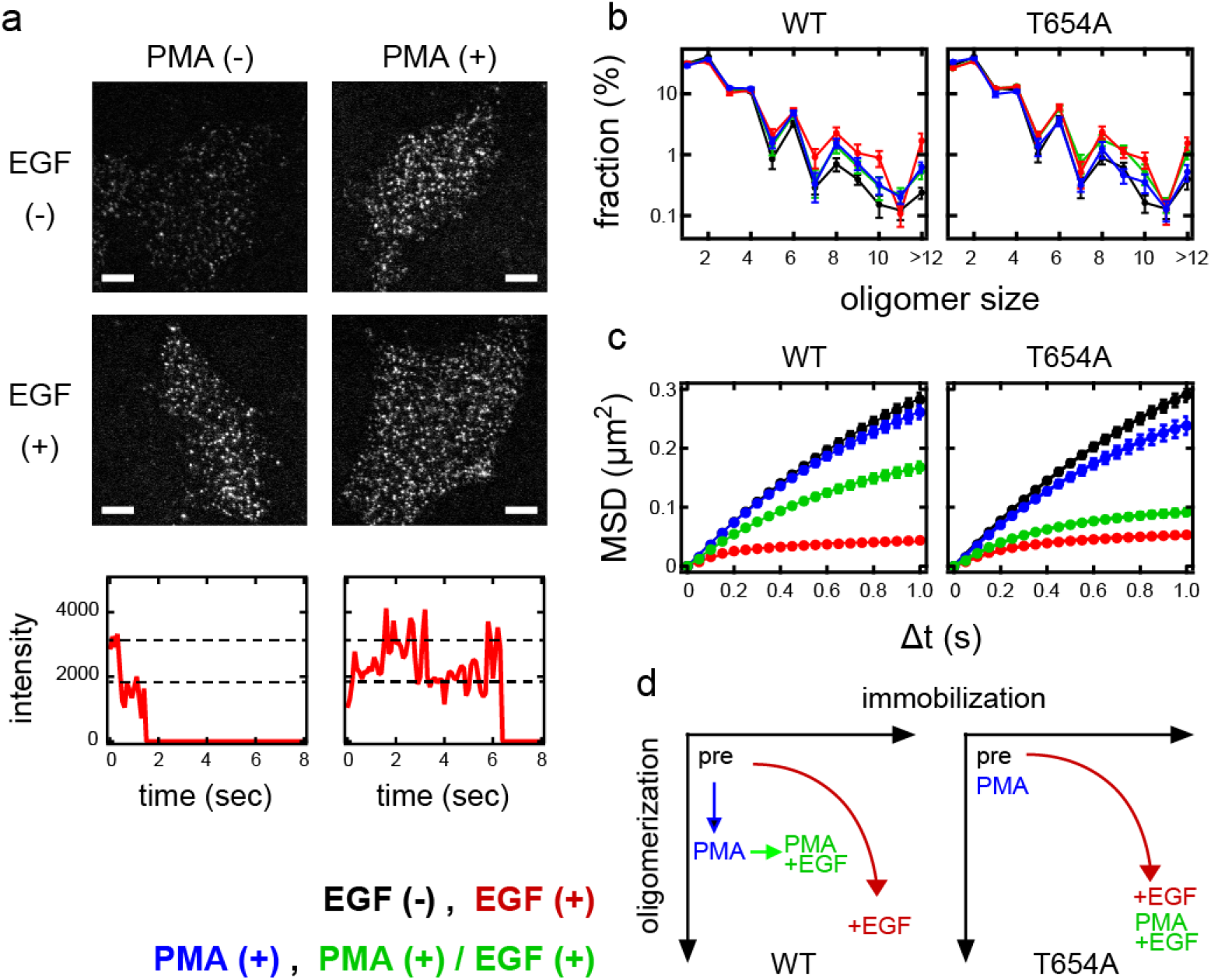
Oligomerization and lateral movements of EGFR on the living cell surface. (**a**) Single molecule imaging of wt EGFR-GFP on the surface of living CHO-K1 cells with (right) and without (left) PMA pretreatment. Cells were stimulated with (lower) and without (upper) EGF. Bar, 5 μm. Quantized transitions in the fluorescence intensities (bottom) indicate single-molecule resolution of the imaging. (**b**) Oligomer size distributions of wt (left) and T654A mutant (right) EGFR in cells. The oligomer size ratio was measured before and 10 min after EGF stimulation. (**c**) Mean square displacement (MSD) of wt (left) and T654A (right) EGFR spots as a function of the time interval, indicating lateral mobility. The MSD was calculated before and 10 min after EGF stimulation. In **(b, c)**, cells were pretreated with (blue, green) or without (black, red) PMA and stimulated (red, green) or not (black, blue) with EGF. (**d**) Diagram of the oligomerization and immobilization states of wt (left) and T654A mutant (right) EGFR suggested from single-molecule measurements. Arrows indicate the state transitions after PMA treatment and EGF stimulation.

Even in the absence of EGF, PMA treatment of cells increased fractions of higher-order wt EGFR oligomers (Fig. 10b left), though diffusion movements were hardly affected (Fig. 10c left). This oligomerization was not as strong as that induced by EGF in the absence of PMA, and application of EGF to the PMA treated cells did not induce further oligomerization at least up to 10 min but significantly decreased mobility. While in cells without PMA treatment, application of EGF induced strong oligomerization and immobilized wt EGFR. For T654A mutant, PMA treatment hardly affected both oligomerization (Fig. 10b right) and movements (Fig. 10c right) in the absence of EGF. EGF induced strong oligomerization and immobilization of T654A mutant independent of PMA treatment. The effects of PMA to induce EGFR oligomerization that was dependent on Thr654 were consistent with pT654-induced oligomerization of TM-JM peptides in nanodiscs.

In summary, single-molecule measurements suggest three states of EGFR oligomerization depends on pT654 and EGF association (Fig. 10d). PMA treatment of wt EGFR but not T654A mutant induced a medial level of oligomerization, which could be stabilized by pT654. Strong oligomerization observed under the weak pT654 level in wt EGFR and no pT654 in T654A mutant was caused by a distinct mechanism of pT654. On the other hand, it is possible that immobilization of EGFR relates with its tyrosine phosphorylation levels as we have observed previously (29). The EGF-induced immobilization in the presence of PMA was more evident in T654A mutant in which pY1068 level was higher than that in wt EGFR (Fig. 9), and PMA treatment decreased immobilization and pY1068 in wt EGFR.

### Interaction of EGFR with GRB2

We finally measured the interaction of EGFR with GRB2 in living cells using a split luciferase (NanoBiT) assay, in which the C-terminus of EGFR and the N-terminus of GRB2 were conjugated with the large BiT (LgBiT) and the small BiT (SmBiT) of NanoLuc luciferase (30), respectively. The association of EGFR and GRB2 promoted the formation of active luciferase to produce chemiluminescence emission (Fig. 11a). In the timecourses of the NanoBiT signal increases after EGF treatment of cells expressing the wt or T654A mutant EGFR (Fig. 11b), the maximum intensities indicated a dose dependent response to the EGF concentration in the medium (Fig. 11b, c). The maximum intensity was significantly increased after pretreatment with PMA in cells expressing wt EGFR, despite the fact that the pY1068 level was reduced after the PMA treatment (Fig. 9c). This effect of PMA was not observed to any extent with the T654A mutant, and the GRB2 association was not increased by this mutation in the absence of PMA, even though the pY1068 level after EGF stimulation was significantly increased in the mutant (Fig. 9c). The increase in GRB2 association observed for wt EGFR after PMA treatment was not likely to be an indirect effect of PMA because it was not observed for the T654A mutant, in which the pY1068 level was also affected by PMA. These results suggest that pT654 promotes the formation of a GRB2 recognition state for EGFR, and that the inhibition of Thr654 phosphorylation prevents a GRB2-EGFR association in spite of enhanced Tyr1068 phosphorylation.

**Figure 11.**
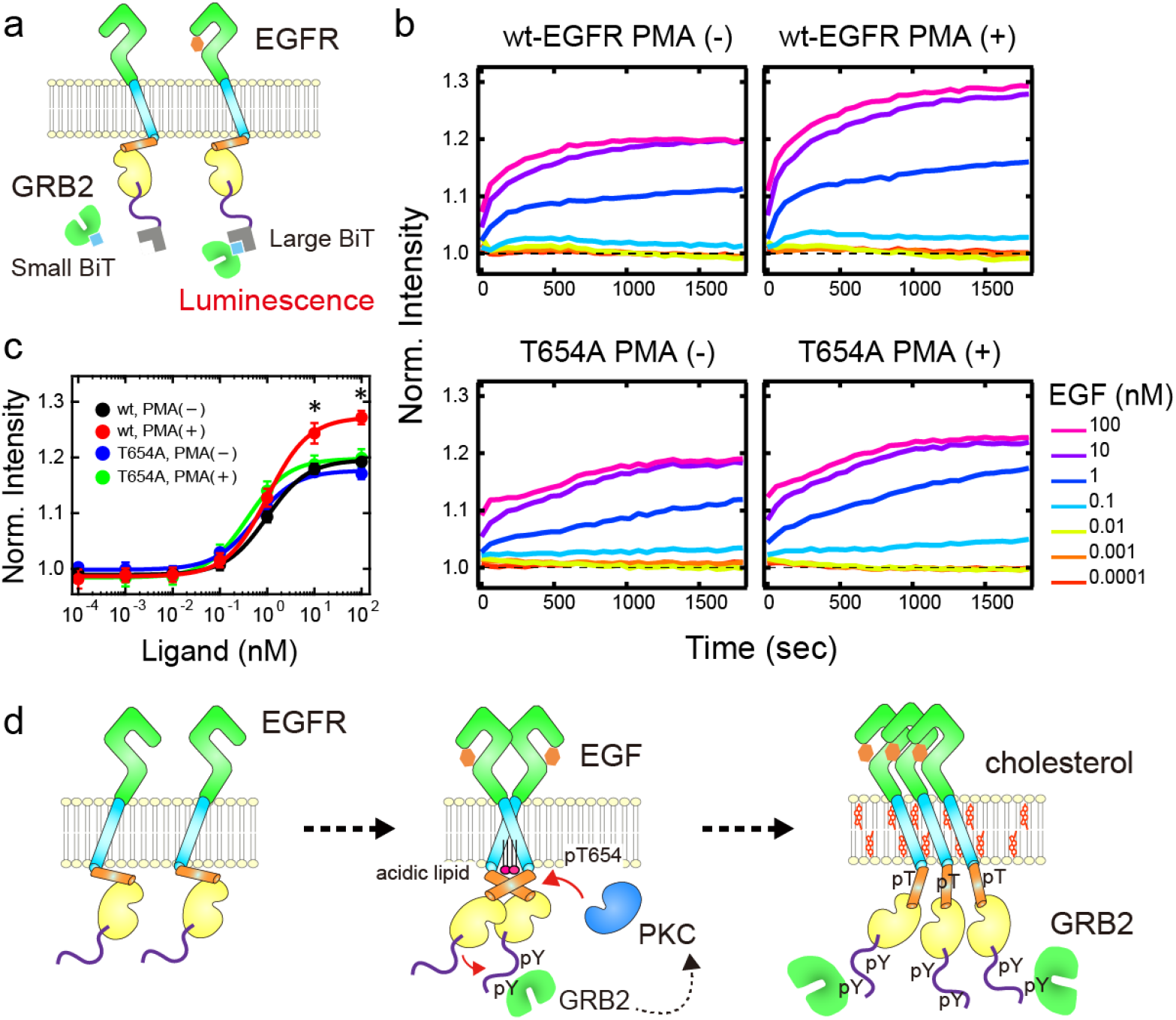
NanoBiT assay for the EGFR/GRB2 interaction in living cells. (**a**) Schematic illustration of the NanoBiT assay of EGFR/GRB2 interactions. (**b**) Typical time courses of chemiluminescence signals generated from the complex formation of large BiT (LgBiT)-fused EGFR and small BiT (SmBiT)-fused GRB2 after EGF application at time 0. The final concentration of EGF in the culture medium was varied from 0.0001 to 100 nM. Measurements were done in cells with (+) or without (-) PMA pretreatment. (**c**) Dose dependency of the chemiluminescence intensities at 30 min after EGF stimulation. The average values from four independent experiments are shown with SE. Lines indicate fitting with a Hill-equation function. *p < 0.05 determined by Etest against the signal in wt cells without PMA. (**d**) A schematic model of the activation and signal transduction process for EGFR dimers and oligomers. In the oligomers of EGFR, affinity with GRB2 is increased.

## Discussion

We have here studied the dimerization and oligomerization of EGFR molecules by reconstituting its TM-JM peptides into nanodiscs. As expected from the positively charged JM-A sequence and accumulation of EGFR in the raft membrane, PS and cholesterol affect the molecular assembly of the TM-JM peptides. Interestingly, these two lipid species each function in a specific fashion i.e. the *E_FRET_* distributions between the two TM-JM peptides in the nanodiscs suggested that PS facilitates JM dimerization, while cholesterol induces closer positioning of both the TM and JM domains (Figs. 3 and 4). In addition, cholesterol promoted the oligomeric assembly of TM-JM peptides (Figs. 6 and 7). We herein propose schematic models for the formation of the EGFR TM-JM dimers and trimers under various conditions of lipid exposure and Thr654 phosphorylation (Fig. 8), in which PS and cholesterol exert competitive effects on dimerization and oligomerization, and pT654 disrupts the PS-induced JM dimer, thus promoting oligomerization of the peptides.

It should be noted that the *E_FRET_* distribution was broad in every one of our observations in this present study, especially between JM domains, indicating multiple configurations coexisting under each condition. Non-phosphorylated peptide dimers showed a peak at around *E_FRET_* > 0.90 and > 0.81 both for N- and C-terminus labeling, respectively. We can attribute this configuration to that suggested in a previous NMR study, in which two JM domains form an anti-parallel helix dimer (9). PS stabilized this configuration probably at the JM side of the non-phosphorylated peptide. Acidic lipids are known to interact with the positively charged JM-A domain (19, 31) whereas cholesterol induced more proximation of the two peptides at the both N- and C-termini in the major configuration (*E_FRET_* 0.95 and 0.93), which must be distinct from the arrangement containing antiparallel JM helices (Fig. 8). If the TM-JM domains of whole EGFR dimer adopt similar configurations as suggested for the TM-JM peptides in the nanodiscs, the arrangement of two kinase domains indicates namely, the kinase activity would be affected by the lipid composition and by pT654 (Fig. 11d).

Cholesterol was found in our current analysis to increase the population of nanodiscs containing three TM-JM peptides with pT654 (Fig. 5). The fluorescence intensity distributions of the two Cy3 probes among the three peptides in the presence of cholesterol suggested that a close trimer was the major configuration (Figs. 6 and 7). The cholesterol-induced oligomerization of TM peptides with a short JM region (to T654) of EGFR in liposomes has been reported previously from NMR analysis (32). In that report however, pT654 in the peptide showed no obvious effect on the oligomerization in PC and PC/cholesterol liposomes (without any acidic lipids), consistent with our current and previous results indicating interplay of acidic lipids and pT654. The induction of oligomerization seems to be a general effect of cholesterol upon α-helix peptides in lipid bilayers (33). Our result does not necessarily mean trimerization in the physiological conditions but reflects a tendency of oligomerization. We picked up trimers in the analysis to avoid complexity in the interactions between more than three molecules. While in the PC/PS membrane without cholesterol (Figs. 6 and 7), the probability to adopt a one dimer + one monomer configuration seems to be increased for the non-phosphorylated peptides, likely because peptides have difficulty forming trimers when containing the anti-parallel helix JM dimer, the pT654 event appears to dissociate the JM dimer to help in the formation of the close trimer especially in the presence of cholesterol. If this assumption is correct, oligomer formation will be inhibitory for EGFR kinase activity. Our observed increases in the pY1068 level in the T654A mutant of EGFR (Fig. 9) support this possibility.

Recently, we studied the effects of cholesterol depletion on EGF signaling under the same experimental condition as used in this study (27). Treatment of cells with methyl-β-cyclodextrin increased the fraction of preformed EGFR dimers and induced hyperphosphorylation of Y1068 and Y1173 of EGFR upon EGF stimulation. Nevertheless, cholesterol depletion diminished reactions downstream of EGFR autophosphorylation, including the translocation of GRB2 to the cell surface and phosphorylations of SHC, AKT, and ERK. It also strongly inhibited the EGF-induced clustering of EGFR over dimers. These observations are consistent with those of the current study, suggesting biphasic functions of membrane cholesterol on the EGF signaling. Cholesterol inhibits EGFR dimerization for kinase activation but promotes oligomerization of the (relatively small amount of) tyrosine-phosphorylated EGFR to enhance signaling to the downstream molecules.

The antiparallel helix dimer of JM is thought to facilitate asymmetric interaction between the kinase domains of EGFR, and hence its activation, in order to phosphorylate tyrosine residues in the C-tail (11, 34). This tyrosine phosphorylation results in the recruitment of PKC and other threonine kinases from the cytoplasm to the EGFR molecules for the phosphorylation of Thr654, which is known to negatively regulate EGFR signaling (26, 35). Our previous results suggested that the mechanism underlying this negative effect of pT654 is the dissociation of JM dimers in the presence of acidic lipids (20). At the same time, pT654 might induce the oligomerization of EGFR in the presence of cholesterol. Supporting this possibility, our current single-molecule imaging in living cells revealed the oligomerization of unliganded wt EGFR after PMA treatment, which induced pT654 (Fig. 10). In the unliganded condition, the extracellular domains of EGFR are thought to prevent the JM dimerization for kinase activation (9, 36). Clustering of EGFR is also prevented largely in the unliganded condition. However, probably due to the structural fluctuation, a small part of the unliganded EGFR oligomerized (27). pT654 (PMA) stabilized the unliganded oligomers possibly destructing the JM antiparallel dimers.

We previously reported that oligomers of EGFR formed after cell stimulation with EGF function as the major signal transduction sites for GRB2, where the dissociation rate constant with GRB2 was reduced increasing the affinity between single-molecules of EGFR and GRB2 (14). In addition, our current analyses found an increase in the wt EGFR/GRB2 association following PMA treatment (Fig. 11). We speculate that after the initial Tyr phosphorylation, successive Thr phosphorylation reduces Tyr kinase activity of EGFR but enhances the formation of signal transduction oligomer in the medium immobilized and oligomerized state of EGFR molecules (Fig. 10d). Further immobilization and oligomerization were found in our current experiments to be induced by EGF in the absence of PMA for wt EGFR and with or without PMA for the T654A mutant. This process might include EGFR molecules accumulated into the clathrin coated pits (37) and be independent of pT654. In the previous report (38), ERK activation was reduced by the Thr654/669 of EGFR to Ala substitution suggesting only negative effects of Thr654, however, oligomerization of EGFR was not observed in this report. In addition, because tyrosine phosphorylation is a prerequisite for EGFR signaling, strong inhibition of the kinase activation will prevent the downstream signaling even after the oligomerization of EGFR. In our experimental condition, even though pT654 was inhibitory for EGFR kinase activity, it promoted signal transduction downstream.

Based on our present results, we propose a model of EGFR signaling regulated by membrane lipids and Thr654 phosphorylation (Fig. 11d). The signal transduction mediated by EGFR is a complex multi-step process. Conformational changes in the extracellular domain of EGFR upon ligand association allow JM domains to form anti-parallel JM helix dimers and asymmetric kinase domain dimers(11, 34). This is the activation process for EGFR kinase, in which acidic membrane lipids and cholesterol play stimulative and inhibitory roles, respectively (20, 27). Tyrosine phosphorylation in the kinase-active EGFR dimers recruits PKC from the cytoplasm (39). The association of PLCγ to the EGFR phosphotyrosine for the degradation of PIP2 is involved in this process. PKC then phosphorylates Thr654 (40), which dissociates anti-parallel JM dimers in the presence of remaining acidic lipids and supports the oligomerization of EGFR in the presence of cholesterol, especially after the removal of acidic lipids around the EGFR molecules. The cholesterol-induced oligomer of EGFR is a major site of interaction with cytoplasmic proteins including GRB2 (14). Thus, a major function of EGFR is shifted from a kinase for self-activation to a scaffold for signal transduction. Thr654 phosphorylation is a key step underlying this role change of EGFR and is not merely an inactivating mechanism. The degradation of PIP2, a major anionic lipid in the inner leaflet of the plasma membrane, may support this role change. Importantly, both Thr654 phosphorylation and PIP2 degradation are caused by the kinase activation of EGFR. Hence, this represents an ingenious autoregulatory process involving membrane proteins and lipids.

## Materials and Methods

### Materials

1-palmitoyl-2-oleoyl-*sn*-phosphatidylcholine (PC), 1-palmitoyl-2-oleoyl-*sn*-phosphatidylserine (PS), and cholesterol were purchased from Avanti Polar Lipids (Alabaster, AL) as chloroform solutions (PC and PS) or powders (cholesterol). Cy3-maleimide and Cy5-maleimide were purchased from GE Healthcare Life Sciences (Little Chalfont, UK). n-octyl-b-D-glucoside (OG) was purchased from Dojindo (Kumamoto, Japan). Monofunctional polyethylene glycol-succinimidyl valerate (s-PEG, 5000 mol wt) and biotinylated monofunctional polyethylene glycol-succinimidyl valerate (b-PEG, 5000 mol wt) were purchased from Laysan Bio (Arab, AL). Chinese hamster ovary K1 (CHO-K1) cells were provided from RIKEN BRC through the National Bio-Resource Project (MEXT, Tokyo, Japan).

### Plasmid construction

Construction of the cDNA of full-length human EGFR (wt) fused with GFP was described previously (14). T654A mutant DNA was constructed using PrimeSTAR Max (Takara, Kusatsu, Japan) in the wt EGFR vector. The primer sequences were as follows: EGFR(T654A)-f: GAAGCGCGCGCTGCGGAGGCTGCTGC and EGFR(T654A)-r: CCGCAGCGCGCGCTTCCGAACGATGTG, respectively. For NanoBiT assays, full-length human EGFR (wt or T654A mutant) was fused with LgBiT at the C-terminus (wt or T654A EGFR-LgBiT), and GRB2 was fused with SmBiT at the N-terminus (GRB2-SmBiT) as follows. The LgBiT fragment amplified from pBiT1.1-C [TK/LgBiT] Vector (Promega) using KOD One PCR Master Mix (TOYOBO) was subcloned into the AgeI- and NotI-digested EGFP-N1 vector (Clontech), and subsequently full-length EGFR fragment was subcloned into the NheI- and HindIII-digested the LgBiT-inserted EGFP-N1 vector. The GRB2-SmBiT fragment was constructed using KOD One PCR Master Mix (TOYOBO), and subcloned into the AgeI- and SalI-digested EGFP-C2 vector (Clontech). The primer sequence of SmBiT was designed from pBiT2.1-N [TK/SmBiT] Vector (Promega).

### Peptide synthesis and purification

Peptides corresponding to the TM-JM regions of EGFR (618-666) were synthesized by solid-phase methods with the sequence KIPSIATGMVGALLLLLVVALGIGLFM-RRRHIVRKRT_654_LRRLLQERELVE-NH_2_ (28). For the experiments with the C-terminal labeled EGFR peptide, peptides containing a cysteine at the C-terminus were synthesized. These synthetic peptides were purified by reverse-phase high-performance liquid chromatography on a C4 column with a gradient of 1-propanol and acetonitrile (1:1) over 0.1% aqueous trifluoroacetic acid. To prepare the C-terminal labeled peptide, Cy3-maleimide or Cy5-maleimide was introduced to the sulfide group on the cysteine at the C-terminus of the TM-JM peptide by mixing the peptide and the fluorescence derivative in dimethylformamide under basic conditions. For experiments with the N-terminal labeled peptide, Cy3-COOH or Cy5-COOH was reacted with an elongating peptide on the resin in the presence of 1-[bis(dimethylamino)methylene]-1H-benzotriazolium 3-oxide hexafluorophosphate (HBTU) and diisopropylethylamine (DIEA), which activate the carboxyl group on the fluorophore derivative. For synthesis of Thr654 phosphorylated peptides, phosphorylated threonine derivatives were utilized. The purity was confirmed by reverse-phase high-performance liquid chromatography and matrix-assisted laser-desorption/ionization time-of-flight mass spectroscopy analysis.

### Nanodisc preparation

For nanodisc construction, fluorescent EGFR TM-JM peptides co-solubilized with lipids and OG in hexafluoroisopropanol were first dried to form thin films. These peptide films were then resolubilized in buffer A (0.5 M NaCl, 20 mM Tris/Cl, 0.5 mM EDTA) containing 30 mM OG and 5 mM dithiothreitol (pH 7.5). His8-tagged MSP 1E3D1 (MSP) was expressed in *E*. coli and purified as described previously (41). The concentration of MSP was quantified based on the absorbance at 280 nm (29,910 M^−1^cm^−1^). Thin PC or PS films were formed by evaporation of the solvent (chloroform) under a steam of nitrogen gas and dried in vacuum. Cholesterol powders were first dissolved in chloroform, and a thin film was formed as described above. PC, PS, and cholesterol were resuspended in buffer A containing 0.4 M sodium cholate (pH 7.5) at a final concentration of 10 mM. Cy3- and Cy5-labeled TM-JM peptides in buffer A were mixed in equal amounts and then conjugated with MSP and phospholipid mixtures (PC, PC/PS, PC/cholesterol, PC/PS/cholesterol) at a molar ratio of 1:1:120 μM (TM-JM/MSP/lipids). The mixture was dialyzed against a buffer containing 0.5 M NaCl, 20 mM Tris/Cl, and 5 mM EDTA (pH 7.5) at 4°C to reconstitute the nanodiscs by removing the detergent. The aggregates and liposomes were removed from the mixture by size-exclusion chromatography using a Superdex 200 Increase column (GE Healthcare Life Sciences) and the peak fractions containing nanodiscs of around 11 nm in diameter were collected.

### Single-pair FRET (spFRET) measurements

Nanodisc samples were immobilized on the surface of a glass chamber as described previously (20, 43, 44). Briefly, amine-modified glass surfaces were coated with 99% s-PEG and 1% b-PEG. NeutrAvidin (Thermo Fisher Scientific, Waltham, MA) was then bound to the b-PEG. The nanodisc samples bound with biotinylated anti-His8-tag antibody (MBL Life Science) were loaded into the glass chamber and allowed to bind to the NeutrAvidin-coated glass surface, after which unbound nanodiscs were washed away. To reduce the photobleaching rate of Cy3 and Cy5, the nanodisc-loaded chamber was filled with dialysis buffer containing 2-mercaptoethanol at the final concentration of 0.5% (w/v). The fluorescence of Cy3 and Cy5 was observed under a TIRF microscope based on an inverted microscope (Ti2; Nikon) with a 60x oil-immersion objective (ApoTIRF 60x 1.49 NA; Nikon). The fluorescence activity of Cy3 was excited using a 532 nm laser (Compass 315M-100). Dual-color imaging was carried out through a 4x relay lens by using two EMCCD cameras (C9100-134, ImagEM; Hamamatsu Photonics, Hamamatsu, Japan) with a 200x EM gain. Images of 512 × 512 pixels (67 nm/pixel) were recorded with a temporal resolution of 100 ms/frame using MetaMorph (Molecular Devices, San Jose, CA) or AIS (ZIDO, Toyonaka, Japan).

### Analysis of FRET signals

The measurement of fluorescence intensities of single nanodiscs was performed using ImageJ software, as described previously (45). The background noise was filtered out using the Subtract Background function in ImageJ. Fluorescence intensities of Cy3 and Cy5 in single nanodiscs were measured as averages from circles with a diameter of 12 pixels containing a fluorescence spot. The average intensity of the same sized circles in which no spot was present was subtracted as the background. Along the fluorescence trajectories of TM-JM-Cy3 and TM-JM-Cy5, the FRET efficiency, *E*_FRET_, for each frame (> 2000) was calculated from the fluorescence intensities in the donor *I*_D_ and FRET *I*_A_ channels as

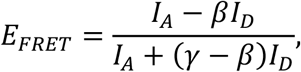

where *β* and *γ* are coefficients for the compensation of fluorescence leakage from the donor dye to the acceptor detector channel, and the difference in the detection efficiencies of the dyes, respectively (46). Coefficients were calculated using the intensity time traces as *β* = 0.03 and *γ* = 0.4, respectively. *E_FRET_* distributions were all different in Kolmogorov-Smirnov (KS) test with a rejection p < 0.006. The peak positions and their 95% percentile section were estimated using bootstrap method from a discretized *E_FRET_* distribution (Table I), in which the bin width was set to 0.02 and the resampling number was 300. KS test was done using “StatsKSTest” in Igor Pro 8.0 (WaveMetrix). Resampling for bootstrap method was done using “sample” function in R.

### Cell culture and transfection

CHO-K1 cells were maintained in HAM F12 medium supplemented with 10% fetal bovine serum at 37°C under 5% CO_2_. HEK293S cells were maintained in DMEM F12 medium supplemented with 10% fatal bovine serum at 37°C under 5% CO_2_. For western blotting assays, DNA constructs of full-length wt and T654A EGFR (1 μg) were transiently transfected into CHO-K1 cells using FuGENE HD Transfection Reagent (Promega, Madison, WI). For single-molecule measurements, CHO-K1 cells were transfected with either a wt or T654A EGFR-GFP gene (0.5 μg each) using Lipofectamine 3000 Reagent (Thermo Fisher Scientific). For NanoBiT assays, HEK293S cells were transfected with a mixture of wt or T654A EGFR-LgBiT gene (1 μg each) and GBR2-SmBiT (0.2 μg) using Lipofectamine 3000 Reagent in 60 mm dish.

### PMA treatment and EGF stimulation

DNA constructs of full-length wt and T654A EGFR were transfected and cultured with 10% fetal bovine serum (FBS) on the day before each measurement. Cells were then starved in modified Eagle’s medium without FBS for 3 hours before the experiment. Phorbol 12-myristate 13-acetate (PMA) was dissolved in DMSO and subsequently diluted in PBS to a final concentration of 10 μM. For PMA pre-treatment, PMA solution was added to the cell cultured medium at a final concentration of 100 nM and incubated for 30 min at room temperature. For EGF stimulation, EGF (PeproTech, Cranbury, NJ) dissolved in PBS was added to the cell cultured medium at a final concentration of 100 ng/mL (for western blotting assays and single-molecule measurements) or as a 0.001 to 100 nM dilution series (for the NanoBiT assay).

### Western blotting analysis

In cells stimulated with EGF for 0, 5, 30 min at 37°C, threonine and tyrosine phosphorylation of the wt and mutant T654A proteins was detected by western blotting using rabbit anti-pT654 (ab75986; Abcam, Cambridge, UK) and rabbit anti-pY1086 antibody (#4407; Cell Signaling Technology, Danvers, MA), respectively. Rabbit anti-EGFR antibody (#sc-03; Santa Cruz Biotechnology, Dallas, TX) was used to detect protein expression. After being resolved by SDS-polyacrylamide gel electrophoresis (PAGE), the electrophoresed proteins were transferred onto a polyvinylidene difluoride (PVDF) membrane and incubated with each antibody (primary antibody) and then with a horseradish peroxidase (HRP)-linked anti-rabbit IgG (secondary antibody; 7076, Cell Signaling Technology). Immunoreactive proteins were detected with Amersham ECL Prime Western Blotting Detection Reagent (GE Healthcare) using an ImageQuant LAS 500 device (GE Healthcare).

### Single-molecule imaging in living cells

The methods for single-molecule measurement and analysis were described elsewhere (Yanagawa and Sako, Methods in Mol Biol, in press; bioRxiv: doi: 10.1101/2020.06.08.141192). The single-molecule imaging of EGFR was performed at the basal plasma membrane of the CHO-K1 cells at 25°C with the same microscopic methods used for the spFRET measurements. The laser wavelength was 488 nm (Sapphire 488; Coherent, Santa Clara, CA) for the excitation of the GFP. Fluorescence images were acquired every 50 ms using AIS software. The acquired multiple TIFF files were processed by ImageJ software as follows: background subtraction was performed with a rolling ball radius of 25 pixels, and two-frame averaging of the images was then performed. Single-molecule tracking analysis was performed with AAS software (ZIDO). All subsequent analyses were performed using smDynamicsAnalyzer (https://github.com/masataka-yanagawa/IgorPro8-smDynamicsAnalyzer), an Igor Pro 8.0-based homemade program.

### NanoBiT assay

HEK293S cells co-transfected with the plasmids of wt or T654A EGFR-LgBiT and GRB2-SmBiT. Overnight after the transfection, cells were collected in 0.5 mM EDTA-containing PBS, centrifuged, and suspended in 2 mL of HBSS containing 0.01 % bovine serum albumin and 5 mM HEPES (pH 7.4) (assay buffer). The cell suspension was dispensed in a 96-well white bottom plate at a volume of 80 μL per well and loaded with 20 μL of 25 μM Nano-Glo Vivazine Live Cell Substrates (Promega) diluted in the assay buffer. After incubation for 2 hrs at room temperature in the assay buffer, cells were pretreated with PMA or vehicle as described above. Basal luminescence was then measured by using a microplate leader (SpectraMax L, Molecular Devices) with an interval of 60 sec at room temperature. After 10 min, 20 μL of the EGF dilution series in the assay buffer or the assay buffer (vehicle) were applied to each well using a benchtop multi-pipetter (EDR-384SR, BioTec, Tokyo, Japan) under red dim light. Then, luminescence was measured for 30 min with an interval of 60 sec. Each time-course of luminescence counts was normalized with the luminescence counts of the vehicle-added well. Dose-response curves were fitted with a Hill-equation to determine the maximum intensity.

## Acknowledgments

YS was supported by MEXT Japan with Grants-in-Aid for Scientific Research (19H05647) and by JST CREST (JPMJCR1912). We thank Hiromi Sato for technical assistance.

## Author contributions

Conceptualization, R.M., Y.S.; methodology, R.M., T.S., M.Y., Y.S.; investigation, R.M., H. T., T. S., M.Y.; manuscript writing, R.M., T. S., Y.S. with feedback from all other coauthors; funding acquisition, Y.S.; supervision, Y.S.

